# Estimation of a Time-varying Apparent Infection Rate from Plant Disease Progress Curves: A Particle Filter Approach

**DOI:** 10.1101/625822

**Authors:** Kaique dos S Alves, Willian B Moraes, Wellington B da Silva, Emerson M Del Ponte

## Abstract

The parameters of the simplest (two-parameter) epidemiological models that best fit plant disease progress curve (DPC) data are the surrogate for initial inoculum (*y*_0_) and the (constant) apparent infection rate (*r*), both being useful for understanding, predicting and comparing epidemics. The assumption that *r* is constant is not reasonable and fluctuations are expected due to systematic changes in factors affecting infection (e.g. weather favorability, host susceptibility, etc.), thus leading to a time-varying *r*, or *r*(*t*). An arrangement of these models (e.g. logistic, monomolecular, etc.) can be used to obtain *r* between two time points, given the disease (*y*) data are available. We evaluated a data assimilation technique, Particle Filter (PF), as an alternative method for estimating *r*(*t*). Synthetic DPC data for a hypothetical polycyclic epidemics were simulated using the logistic differential equation for scenarios that combined five patterns of *r*(*t*) (constant, increasing, decreasing, random or sinusoidal); five increasing time assessment interval (Δ*t* = 1, 3, 5, 7 or 9 time units - t.u.); and two levels of noise (α = 0.1 or 0.25) assigned to *y*(*t*). The analyses of 50 simulated 60-t.u. DPCs showed that the errors of PF-derived 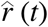 were lower (RMSE < 0.05) for Δ*t* < 5 t.u. and least affected by the presence of noise in the measure compared with the logit-derived *r*(*t*). The ability to more accurately estimate *r*(*t*) using the novel method may be useful to increase knowledge of field epidemics and identify within-season drivers that may explain *r*(*t*) behaviour.

## Introduction

Mathematical models are powerful tools for describing, analysing and comparing plant disease progress curve (DPC) data. In general, these models are derived from two approaches: 1) statistically, or data-driven, when mathematical (population dynamics) models are fitted to observed or simulated DPC data; and 2) fundamentally, or concept-driven, by linking differential equations for describing system behaviour and predicting DPC data (Madden et al. 2007). The fit of complex epidemiological models with multiple parameters is generally challenging as it requires estimation of infection rate (*r*), incubation period, sporulation rate, host growth rate, among others. Hence, simpler models with fewer, yet biologically-relevant, parameters have been preferred and used extensively for describing and comparing plant disease epidemics (Kranz 2003; Madden et al. 2007; Jeger et al. 2011a; 2011b).

Traditionally, and most commonly, two-parameter non-flexible population dynamics models (monomolecular, logistic or Gompertz) are fitted to DPC data,and although more flexible ones, with an additional parameter, are available they are not often used (Berger 1980; Campbell and Madden 1990; Madden et al. 2007). These two parameters describe and allows to predict and compare epidemics: the intercept parameter, representing the disease intensity (*y*) level at time (*t*) zero (*y*_0_), is a surrogate for initial inoculum, and the slope parameter representing the apparent infection rate (*r*), describe the increase in disease intensity per unit of disease intensity per unit of time (Vanderplank 1963; Campbell and Madden 1990). The term “apparent infection rate” refers to appearance of symptoms lagging behind actual infections. Another kind of rate obtained directly from the DPCs, without model fitting, is the absolute rate of disease increase, or the difference (Δ) between disease intensity at two time points, given by 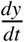. These two rates can be used to compare treatments and identify predictors of epidemic progress. Although it seems appealing to use the absolute rate of disease increase, this rate is inherently much more variable during the course of the epidemics than *r*, not exactly depending on the predictors of its behavior (Berger 1981; Madden et al. 2007). On the other hand, *r* behaves differently: if the epidemics are not disturbed (by a driver), *r* remains constant over time.

The (constant) rate parameter is obtained from fitting models to DPC data over the course of the epidemics. However, as discussed by Madden et al. (2007), it is not reasonable to assume a constant behavior for *r* and fluctuations are expected due to systematic changes in the drivers of infection (e.g. weather favorability, host susceptibility, etc.), thus leading to a time-varying *r* (Madden et al. 2007). In fact, once it is know how *r* changes systematically (increasing, decreasing, oscillating, etc.), modifications of the population dynamics models (logistic, monomolecular, etc.) are possible to account for *r*(*t*) depending on its actual form, which may vary considerably across pathosystems, thus limiting generalizations (Madden et al. 2007).

The traditional way of calculating *r*(*t*) during epidemics was early presented by Vanderplank (1963) and further modifications were proposed 20 years later to correct for host growth (Kushalappa 1980; 1982). However, the uncertainty of the measures of disease intensity at each time point are not handled directly and it is not known how accurate they represent actual *r*(*t*), which cannot be easily measured like disease intensity. In this study we evaluated an alternative method to estimate *r* within two assessment times during the course of the epidemics using one of the various data assimilation modeling techniques. In particular, we tested Particle Filter (PF), a Bayesian method that allows estimating the parameter using measure data with a considerable level of imputed noise (Turner and Sherlock 2013). In brief, the PF algorithm uses a measured state-variable (e.g. *y*) to estimate a parameter/variable (e.g. *r*) at the same time point, which cannot be measured directly. The basic idea behind PF is to approximate the posterior probability of the states using a large number of samples (particles) with their associated weights. These particles and weights are then updated sequentially along with the state evolution when new measures become available (Chen 2003). Primarily used by engineers for solving problems in physics and chemistry, particle filter has been used in human epidemiology to forecast vector-borne epidemics such as influenza (Sheinson et al. 2014; Yang et al. 2014; Dawson et al. 2015; Moss et al., 2016; Ristic and Dawson 2016).

The first application of a filter technique in plant disease epidemiology was the Kalman filter, a predecessor of the PF, used a decade ago for modelling the spatial relationships between grain yield and reflectance measures of wheat streak intensity estimated across the field (Workneh et al. 2009). The authors showed that filters assuming stochastic trends without slopes or deterministic trends with or without slopes, best described yield-reflectance relationships across the field. Nonetheless, no further study in plant pathology have use particle filters as far as known. Our main objective was to evaluate the particle filter as a method for estimating *r*(*t*) from disease progress curve data. We evaluated the method by creating and analyzing synthetic epidemics using a differential form of the logistic model for various scenarios that combined different patterns of simulated *r*(*t*), different interval length between two disease assessment times and two levels of noise in disease data.

## Materials and methods

A common practice when evaluating the performance of a Particle Filter for parameter estimation is the generation of synthetic parameter data (da Costa et al. 2018; Salardani et al. 2018; Dias et al. 2017). Because *r* is unknown (unmeasured) in plant disease epidemics, and assuming that several different forms of *r*(*t*) exist for all sorts of plant disease epidemics, we generated synthetic *r*(*t*) assuming five different patterns: constant, increasing, decreasing, random and sinusoidal. DPCs were then simulated using the differential form of the logistic model, which is suited to describe the behaviour of polycyclic epidemics (Berger 1981).

The use of a known parameter to predict the measures is called “direct problem”, while the “inverse problem” is to use the known (measured/observed) measure to estimate the parameter (Colaço et al. 2012; Kaipio and Somersalo 2005). The inverse problem approach may suffer from inverse crime, or when unrealistic results arise from using the same mathematical model to obtain the predicted and estimated data (Kaipio and Somersalo 2005). To alleviate the issue, discrepancies are introduced, including: 1) adding a noise to the the measured data; 2) using different meshes in the forward and the inverse procedures; or 3) using different mathematical model for the forward and the inverse procedures. In our study, we opted for the first approach and assigned a randomly distributed noise to disease value at the each time point (Kaipio and Somersalo 2005).

### Simulations of *r*(*t*)

We used the differential logistic model to generate DPC data for a hypothetical polycyclic epidemics. Five different patterns of *r*(*t*) were simulated. Initial values and equations varied according to the desired pattern (Table 1). The initial inoculum parameter at time 1 was set to 0.1% disease intensity (*y*_1_ = 0.001) and *r*(*t*) values specific to a time-varying pattern (Equation 1).

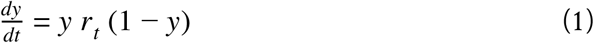

**Table 1:** Initial values of the apparent infection rate (*r*) at time 1 (*r*_1_) and the respective equations for updating *r* (i.e. *r*_*t*+1_) following different temporal patterns, over the course of a simulated polycyclic epidemics of 60 time units using the differential logistic model.

| Time-varying $r$ | Initial value ($r_1$) | Updated value ($r_{t+1}$) |
| --- | --- | --- |
| Non variable (constant) | 0.2 | $r_{t+1} = r_t$ |
| Increasing | 0.05 | $r_{t+1} = r_t + 0.0035$ |
| Decreasing | 0.3 | $r_{t+1} = r_t - 0.0035$ |
| Sinusoidal | 0.2 | $r_{t+1} = r_t + \frac{\pi \cos(\pi \frac{t-1}{30})}{360}$ |
| Random<sup>1</sup> | 0.2 | $r_{t+1} = r_t + 0.05 v_t$ |

<sup>1</sup> $v_t$ is a noise added to rate based on a uniform distribution bounded between -1 and 1.

### Simulations of disease progress curves

We simulated DPCs with five increasing fixed time interval length between two disease assessments (Δ*t* = 1, 3, 5, 7 or 9 time units), meaning that *t* varied from 1 to *N* by Δ*t*, being *N* the duration of the epidemics set to 60 time units. A normally distributed random noise was assigned to predictions of *y*_*t*_ by:

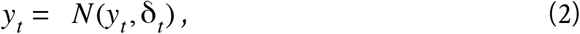

where *N* (*y*_*t*_, δ_*t*_) is normally distributed with mean *y*_*t*_ and standard deviation δ_*t*_ = α*y*_*t*_ (1 − *y*_*t*_), which is non-linear during the epidemic and maximal when disease intensity values reach 50%. We disturbed the epidemics by adding two values of α (noise), 0.1 and 0.25 to *y*(*t*). The ordinary differential equations (ODEs) were solved using the ode() function of the deSolve package (Soetaert et al. 2010) of R (R Core Team 2018).

### Particle filter estimation

Particle Filter (PF) is a Monte Carlo procedure also known as Sequential Monte Carlo (SMC). The main advantage of PF is the combination of the Monte Carlo sampling method and the Bayesian inference, which allows to improve estimation at a low computational cost (Chen 2003). Furthermore, it enables parameter estimation even when considerable noise is associated with the measures. This filter allows to handle nonlinear models, something that the Kalman Filter, its predecessor, could not handle (Chen 2003). We used one of the variants of the PF, the PF by sequential importance resampling (SIR-PF) (Gordon et al. 1993). The SIR-PF includes an additional step (resampling) to avoid particle degeneration, or when increasing weights of variance over time lead to allocation of non-significant weights for particles with the state evolution. This has been a limitation of the sequential importance sampling (SIS) algorithm (SIS-PF), the SIR-PF predecessor (More details in: Gordon et al. 1993, Chen 2003, Colaço et al. 2012, Dias et al. 2017).

In our simulations, the measures were the simulated values of *y* and the parameter to estimate was 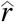, which are linked by an epidemiological model. In the algorithm, both 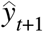 and 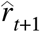 are estimated simultaneously from an independent randomly-generated samples (the particles) of *y*_*t*+1_ and *r*_*t*+1_ obtained from the *priori* 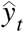 and 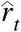.

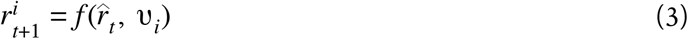

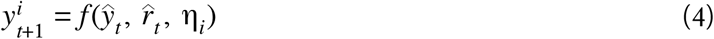

Where *i* = 1, 2, …, *P*, and *P* is the number of particles in the sample, υ_*i*_ is the uniformly distributed noise, and η_*i*_ is the noise arising from a Gaussian distribution. The particles are weighted by their likelihood to the measure *z*_*k*+1_.

Equation 5 provides the weight 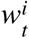 for the respective particle. In Equation 6, the likelihood function, where *W* is the inverse of the covariance matrix, in that, *W* = σ_0_*Inv*_*nxn*_, been σ_0_ a measure of uncertainty (Ozisik and Orlande 2000).

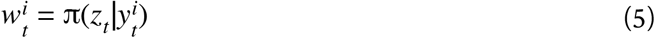

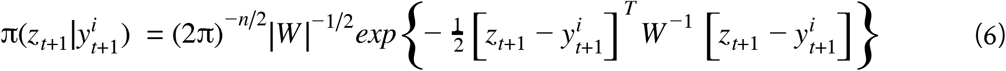

Note that in the likelihood function only *y* is assigned, meaning that the weights of the particles of *y* and *r* are the same. Additionally, Equation 7 provides the weights standardized between zero and one 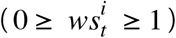. Then, the *posteriori* estimates 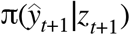 and 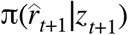 are calculated by a weighted mean (see Fig. S1) of the particles and its respective particle as follows:

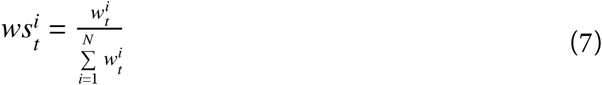

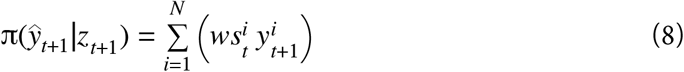

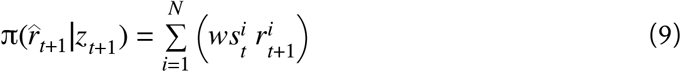

### Setup for the obtaining PF-derived *r*(*t*)

For generating the particles, a 0.5% uncertainty in *y* was assumed (Equation 10).

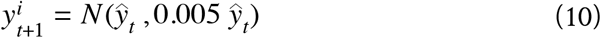

To calculate the weights using the likelihood equation, an uncertainty of 25% was assigned to the measures at each time step of the epidemics. In simulated epidemics, the level of uncertainty assigned to update of the parameter (particle generation for the parameter) was 15% of the previous *r*(*t*) (Equation 11). Therefore, the evolution model for particle generation of y and *r*(*t*) depends on the uncertainty of both in the model and in the parameter, respectively, and is given by

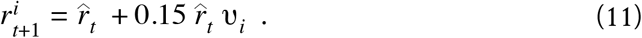

Eventually, the parameter can be updated at each time step based on estimates obtained using another model (weather-based model, for example), but in this work the updates in *r* were assumed to originate from a random process. All estimations were obtained from generating 1,000 particles each time step. The 99% confidence interval (99%*CI*) of *r*(*t*) estimations was given by

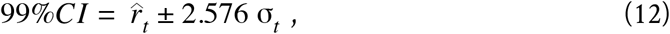

where σ_*t*_ is the standard error of the particles. The error of the estimates of *r*(*t*) was summarized by the root mean square error (RMSE) for simulated values until the epidemics reached 100% severity.

### Apparent infection rate calculation

A reference method for calculating *r*(*t*) was based on the difference in the logits (assuming the logistic model) of two consecutive disease intensity, for each Δ*t* defined previously, divided by the respective length in time units (Equation 15) (Madden et al. 2007).

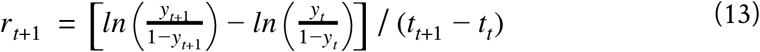

### Data availability and reproducibility

All data from our simulations, analysis and graphical work were conducted using R version 3.5.2 (R Core Team 2018). To fully reproduce our work, a research compendium was organized and all R scripts are available for download (https://osf.io/7nyuj/) (Alves and Del Ponte 2019).

## Results

A total of 50 DPCs were generated from combining five temporal patterns of *r*(*t*), five Δ*t* (time interval) and two α (noise) (Fig. S1). The typical sigmoid-shaped curves were produced for most epidemics. Epidemic curves for Δ = 3 time units, chosen as representative of all patterns produced for the other Δ*t* (Fig S1) are depicted in Fig. 2. All but the DPCs resulted from decreasing *r*(*t*) reached the upper limit (*y* = 1.0), but varied in their shapes and, consequently, in area under the disease progress curves (data not shown). For example, DPCs resulting from increasing *r*(*t*) lagged behind the other DPCs (Fig. 2). The ones resulting from random *r*(*t*) lagged at the beginning of the epidemics but peaked up later in time, reaching maximum earlier than the constant *r*(*t*). The effect of the two α on the shape of the curves was more apparent for the shorter rather than the longer Δ*t* (Fig. S1).

**Fig. 1.**
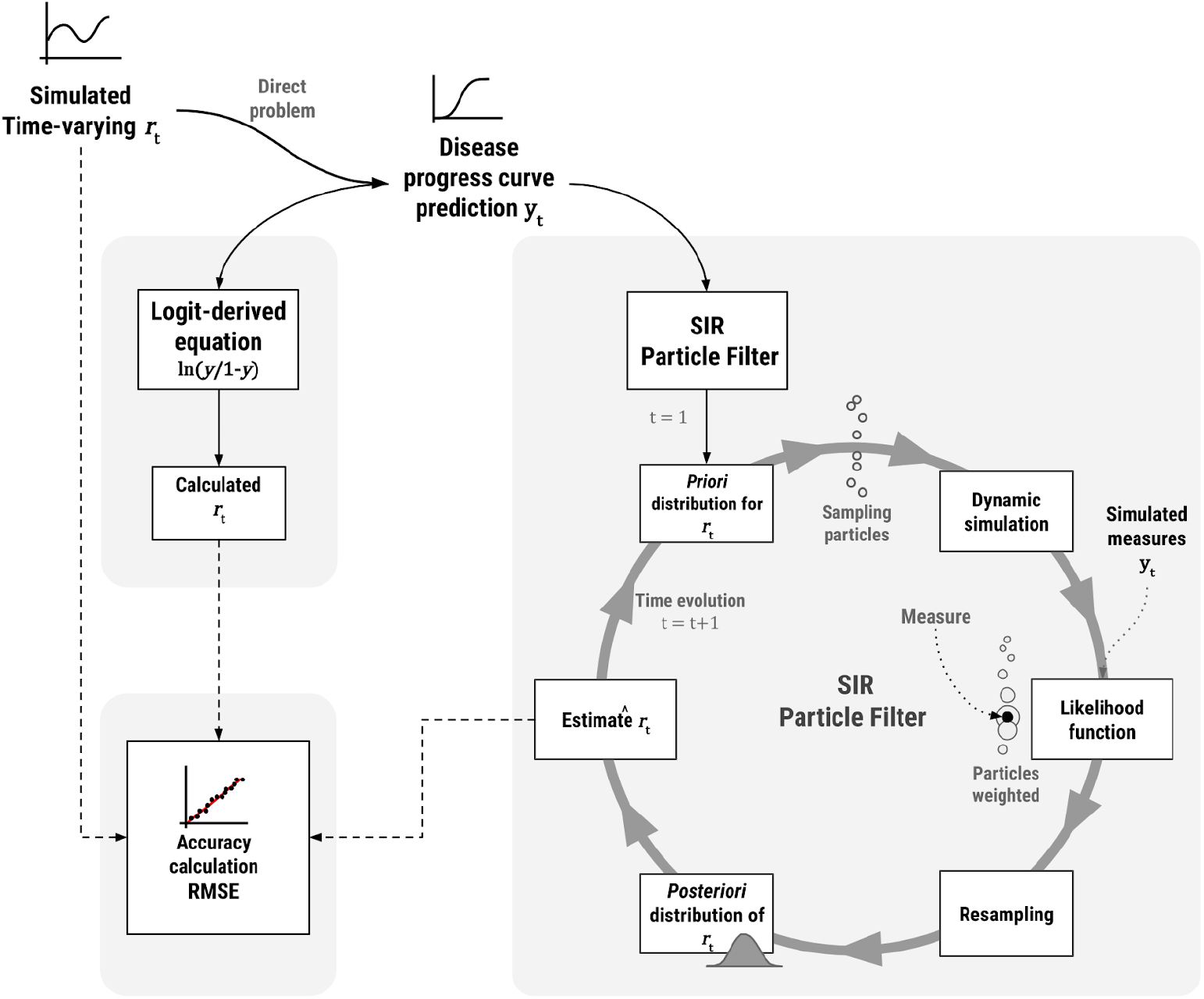
Workflow of the approach for generating synthetic epidemics by the differential logistic model with different patterns of time-varying apparent infection rate - *r*(*t*), and evaluating the performance of a Sequential Importance Resampling Particle Filter (SIR-PF) algorithm to obtain/update 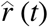 over time. The error of the PF-derived estimates and logit-derived calculated rates were evaluated by the root mean square error (RMSE) as a measure of accuracy. SIF-PF scheme adapted from Dias et al. (2017).

**Fig. 2.**
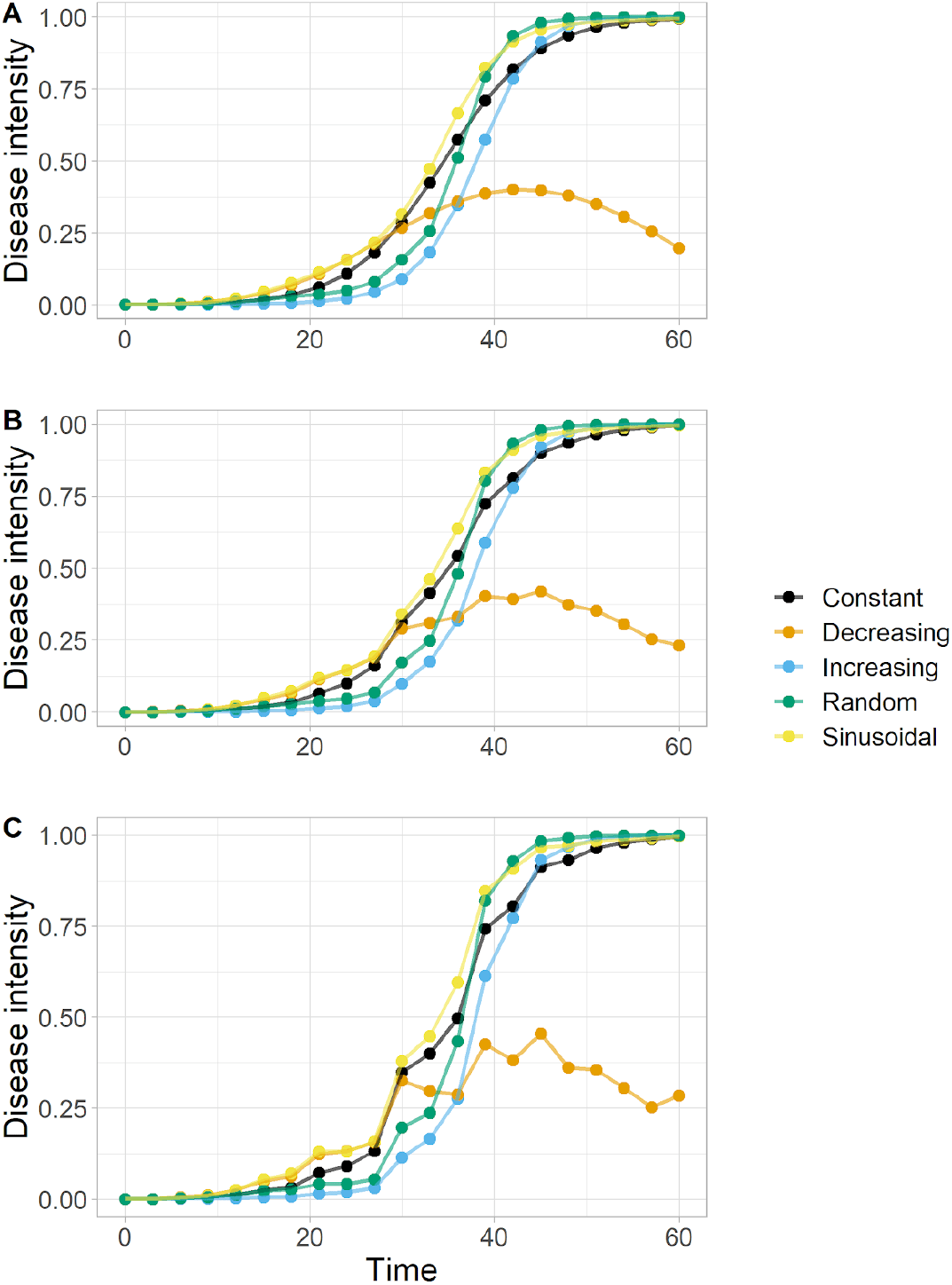
Simulation of disease progress curves (DPCs) by a differential logistic model and five temporal patterns of *r*(*t*) (See equations in Table 1). (**A**) No noise in disease intensity; (**B**) 10% noise (α = 0.1); and (**C**) 25% noise (α = 0.25).

The 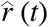 - particle-filter estimates of time-varying rate were generally closer to the synthetic *r*(*t*) than the logit-derived *r*(*t*), regardless of the temporal pattern. However, the superiority of the particle filter compared with logit-derived *r*(*t*) was more evident for scenarios of shortest Δ*t* and largest α than for scenarios of largest-intervals and lowest-noise (Fig. 3, Fig. 4, Fig. S2 and Fig. S3). The two approaches performed similarly when Δ*t* > 5 t.u., especially for the lowest α (10% noise), and > 7 t.u. for the high α (25% noise) (Fig. 3). The SIR-PF allowed to obtain estimates of disease intensity at values that matched quite well the simulated disease intensity which fell within the 99% confidence intervals of the estimates (Fig. S4).

**Fig. 3.**
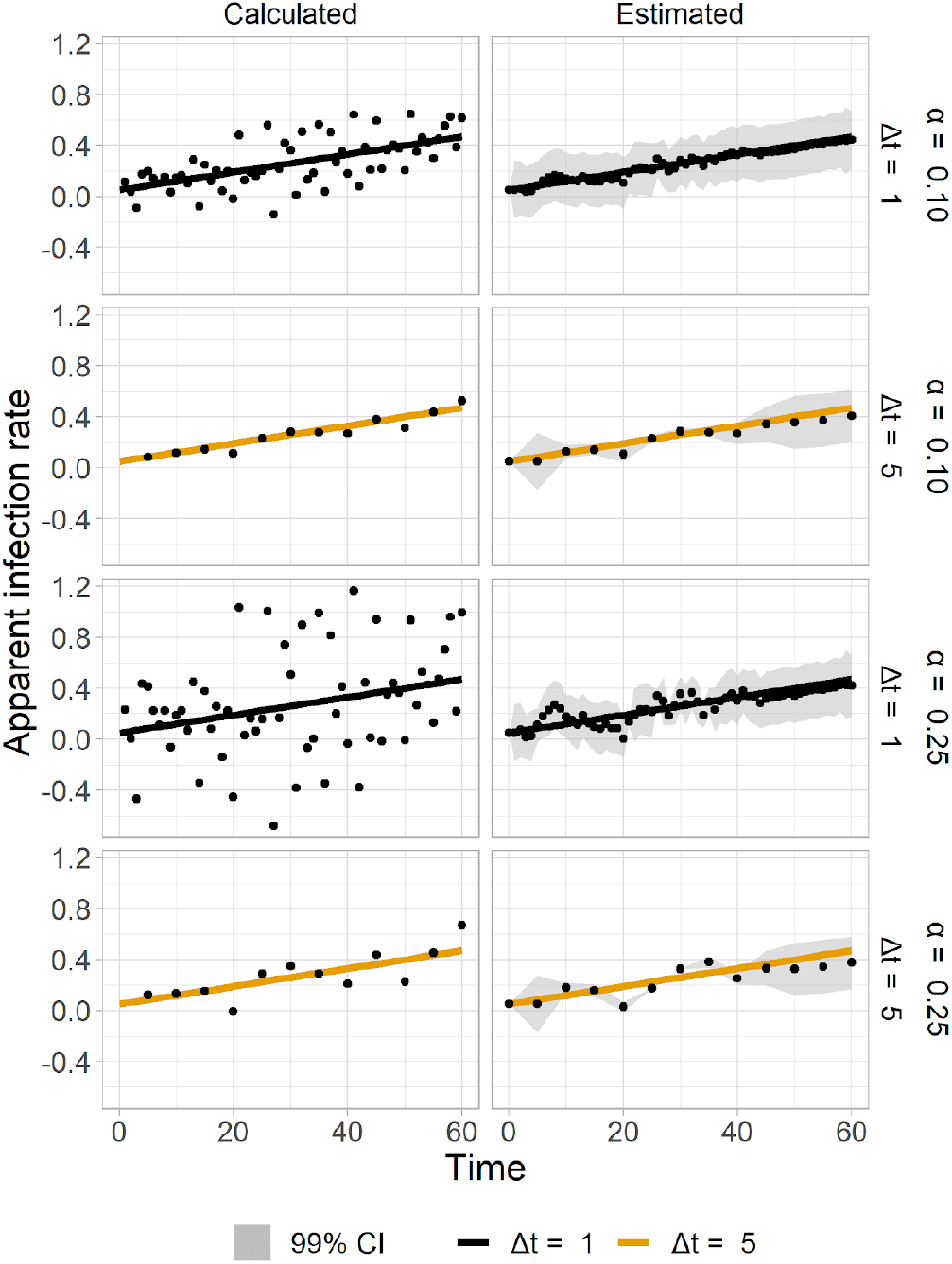
Apparent infection rate *r*(*t*) obtained by a rearrangement of the logistic model (left panel) or an estimation method based on the sequential importance resampling particle-filter approach (SIR-PF) (right panel) for scenarios of two intervals between two assessments (Δ*t* = 1 time unit and Δ*t* = 5 time units) and two levels of noise (α = 0.10 and α = 0.25) assigned to disease intensity data. Solid line represents simulated *r*(*t*) values and particle filter- and logit-derived *r*(*t*) are represented by the dots at the respective panel.

**Fig. 4.**
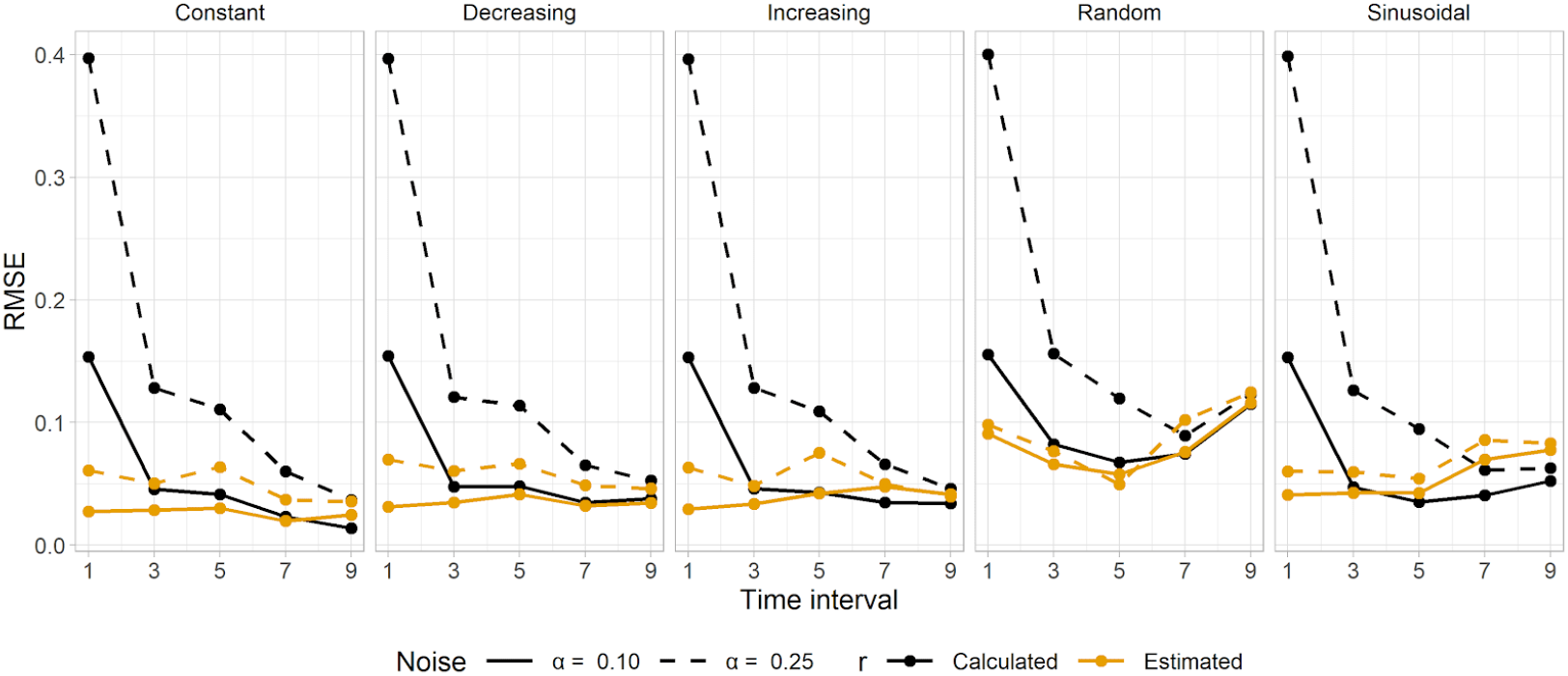
Magnitude of the errors based on a quadratic scoring rule for measuring average error (RMSE) of two methods for recovering the time varying apparent infection rate *r*(*t*), the first a sequential importance resampling particle-filter (SIR-PF) used to estimate *r*(*t*), and the second based on the arrangement of the logistic model, both compared with simulated rates to account for five systematic patterns (each plot in the panel). The methods were evaluated for different time length (in time units) between two assessments and two levels of noise level assigned to simulated disease intensity simulated by a differential logistic model.

## Discussion

In this study, a novel method was proposed and evaluated for estimating the time-varying apparent infection rate *r*(*t*) over the course of synthetic epidemics simulated with variable *r*(*t*) following systematic changes. We showed that the Particle Filter method allowed to obtain estimates of *r*(*t*) with lower error than a simple calculation when interval between time assessment was no greater than five time units and the level of uncertainty of the measures was as high as 25%.

The need to account for time-varying rate in epidemics has been highlighted as one of the challenges in modelling plant diseases (Cunniffe et al., 2015) and solutions to account for a time-varying rate in population dynamics models have been proposed (Madden et al. 2007). Hence, the ability to obtain more accurate estimates of *r*(*t*), especially when disease data are available at relatively short interval (monthly assessment for diseases of perennial crops, for example) between two disease assessments may help to clarify how *r*(*t*) behaves and investigate factors that drive its behavior.

It is well known that the dynamics of field epidemics are disturbed by within-season continuous variations in weather at short (hourly) time scales which affect processes of the disease cycle, mainly the apparent infection rate. For instance, a decline in *r*(*t*) obtained from three assessments in time in Cercospora leaf spot (*Cercospora beticola*) epidemics in sugar beet was associated with variations of temperature that turned less favorable for new infections later in the season (Pundhir and Mukhopadhyay, 1987). Waggoner (1986) assembled mathematical equations to account for parabolic and sinusoidal patterns around an optimum *r*(*t*) in order to reflect the effect of climatic variations on late blight epidemics. Variations in host susceptibility are also known to affect *r*(*t*) and consequently the patterns of the DPCs. An exponential decrease in infection efficiency was observed with leaf aging of *Vitis vinifera* affected by powdery mildew (*Erysiphe necator*) (Calonnec et al., 2018). The same leaf aging effect slowing down epidemics has been reported for powdery mildew in strawberry (Carisse and Bouchard 2010) and myrtle rust in *Eucalyptus grandis* (Xavier et al. 2015). Once drivers are identified, infection rate can be corrected/updated at each time point, which is a common step in process-based simulation models. For example, modifications of a generic rice disease simulation model to account for decreasing or increasing susceptibility of the crop to the disease over time, among other disease-specific parameters, successfully reproduced the typical epidemic curves of rice blast and rice brown spot, respectively (Savary et al. 2012). These curves are somewhat similar to those we produced using the increasing and decreasing *r*(*t*) (Fig. S1).

Our results showed that accuracy the logit-derived *r*(*t*) values were the poorest for scenarios of high noise (uncertainty) and time intervals shorter than 5 time units, a situation in which the particle filter performed better. In fact, particle filter has shown to perform well when recovering information from data even with large signal-to-noise ratio (Dias et al. 2017; Leung et al. 2016; Gordon et al. 1993). These results may be generalized to any time scale (day, week, month, year) and so the advantage of PF becomes more evident. While epidemics in annual crops are usually assessed at a 7 to 10-day interval, epidemics in perennial crops are usually assessed monthly or weekly.

Data assimilation is a well developed technique that has found practical application in other fields and could be useful to tackle other plant pathology problems, as shown previously for a spatial analysis study (Workneh et al. 2009). We showed that an unmeasured parameter (infection rate) can be estimated (recovered) from measured ones (disease intensity), an inverse problem, with higher accuracy than a formula based on the rearrangement of the logistic model which was used to create the synthetic epidemics. We expect that the method may perform similarly for other patterns of *r*(*t*), as well as for other population dynamics models, including the Gompertz (Alves, *unpublished*).

In some situations, a transient state in disease may occur, i.e. momentaneous increase followed by a sudden decrease or vice versa. This is true for some pathosystems where the host is susceptible only for a short time such as Fusarium head blight in small grains - the host is largely vulnerable to infection during relatively short period, restricted to flowering, becoming much less susceptible afterwards (Shah et al. 2018; Del Ponte et al. 2005). In managed epidemics using fungicide spray programs, *r*(*t*) can be greatly reduced but for a relative short time. A recent study reported an effective period of fungicides (inferred from reduction of infection rate) for three sole fungicides and a pre-mix for controlling septoria leaf spot in winter wheat, ranged from 16 to 22 days. However, differences between two absolute rates (treatment and control) were calculated over time to determine when they were equivalent (end of effective protection) (Greiner et al. 2019). The approach developed here could be used to obtain estimates of *r*(*t*) with the advantage of accounting for measures of uncertainty.

The possibility to obtain more realistic estimates of *r*(*t*) may help to increase our understanding of the epidemics and identify within-season drivers of the *r*(*t*) behaviour, which could incorporate disease prediction models. Further studies on this specific problem should focus on using data from actual field epidemics. If an informative model (e.g. weather-driven) for *r*(*t*) is developed, future disease levels can be predicted using real-time or forecast weather. Further, besides estimation of an epidemiological parameter, an unmeasured state variable could be estimated such as the number of plants at the asymptomatic stage for diseases of relatively long incubation period and for which an informative weather-driven model (oscillating temperature) could be useful as well.

## Acknowledgements

The authors are thankful to the Programa de Pós-graduação em Fitopatologia da Universidade Federal de Viçosa and CNPq - Conselho Nacional de Desenvolvimento Científico e Tecnológico for providing a graduate scholarship to K. S. Alves and the CNPq for a research fellowship to E. M. Del Ponte.

## Supplemental material

**Fig. S1.**
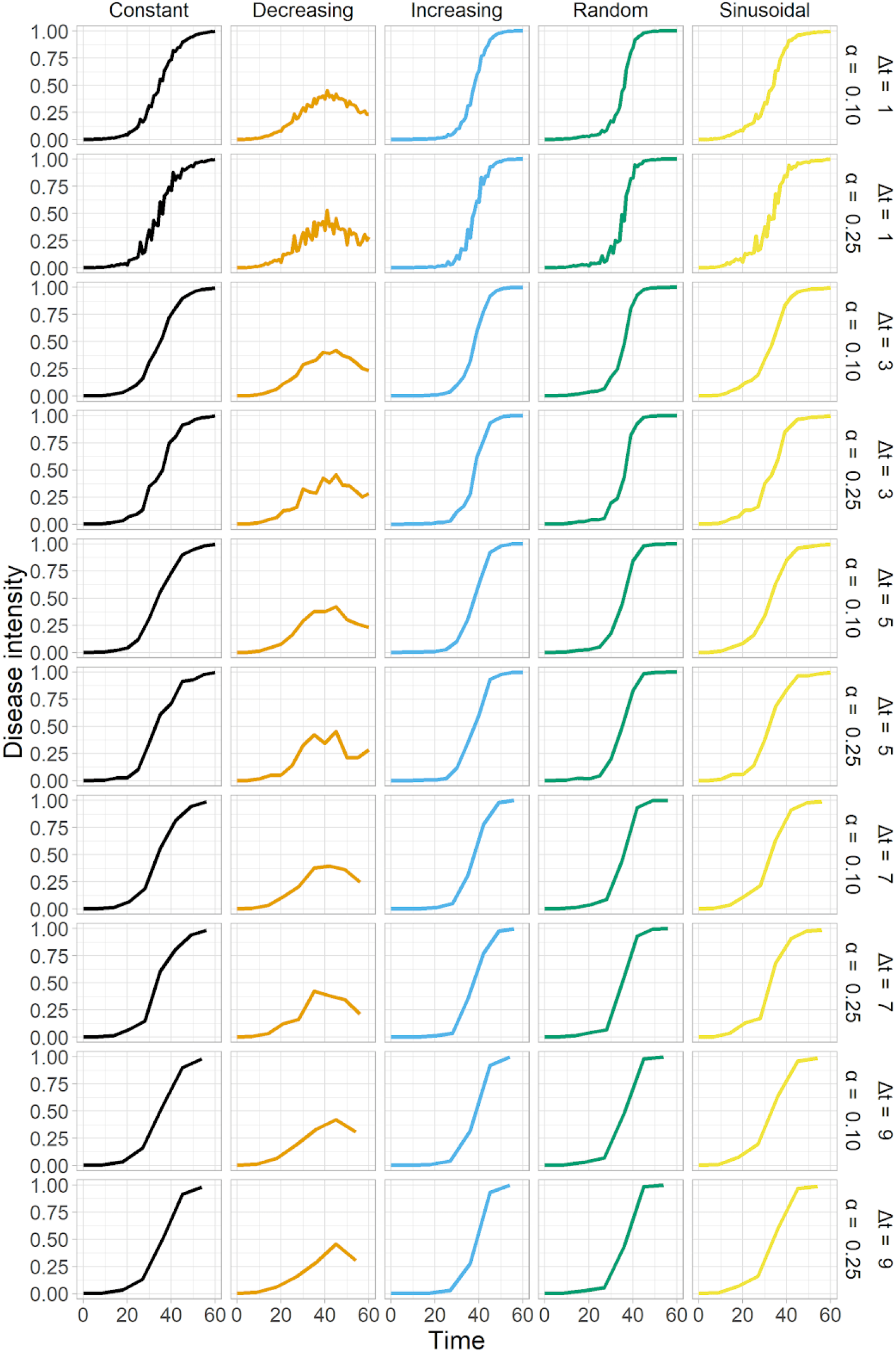
Disease progress curves of 60-day epidemics simulated by a differential form of the logistic model for five patterns of time-varying apparent infection rate *r*(*t*) (constant, decreasing, increasing, random and sinusoidal), five intervals in time units (Δ*t* = 1, 3, 5, 7 and 9) and two levels of noise (α = 0.1 and 0.25) assigned to disease intensity data.

**Fig. S2.**
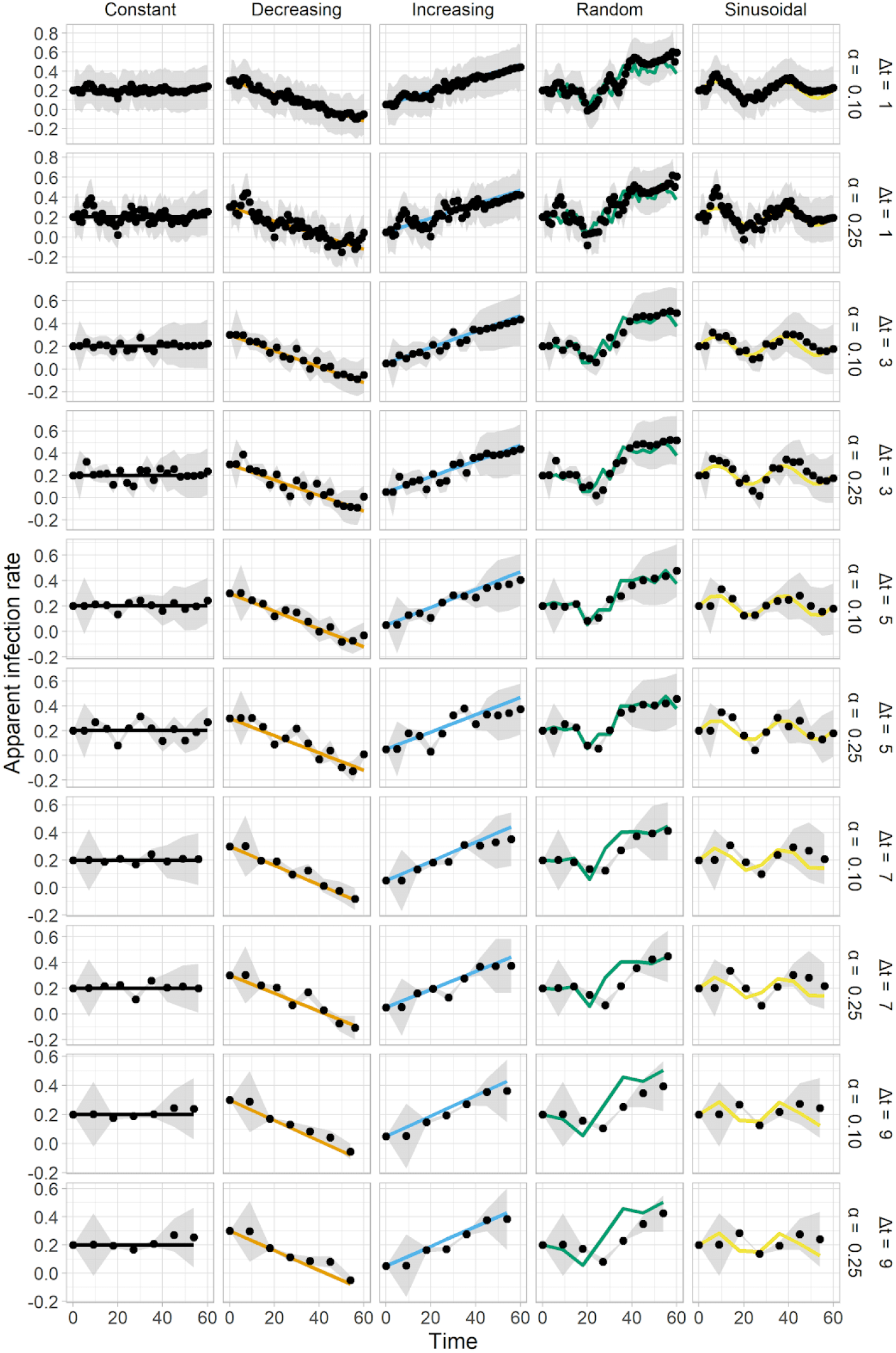
Particle filter estimates of a time-varying apparent infection rate in synthetic disease progress curves simulated by a logistic model with five predefined patterns of time-varying apparent infection rate *r*(*t*) (constant, decreasing, increasing, random and sinusoidal), five intervals in time units (Δ*t* = 1, 3, 5, 7 or 9) and two levels of noise (α = 0.1 or 0.25) assigned to disease intensity data. Solid dots and triangles represent the estimates for the respective level of noise and solid lines represents simulated values of *r*(*t*). The shaded area represents the 99% confidence interval of the particle-filter estimates.

**Fig. S3.**
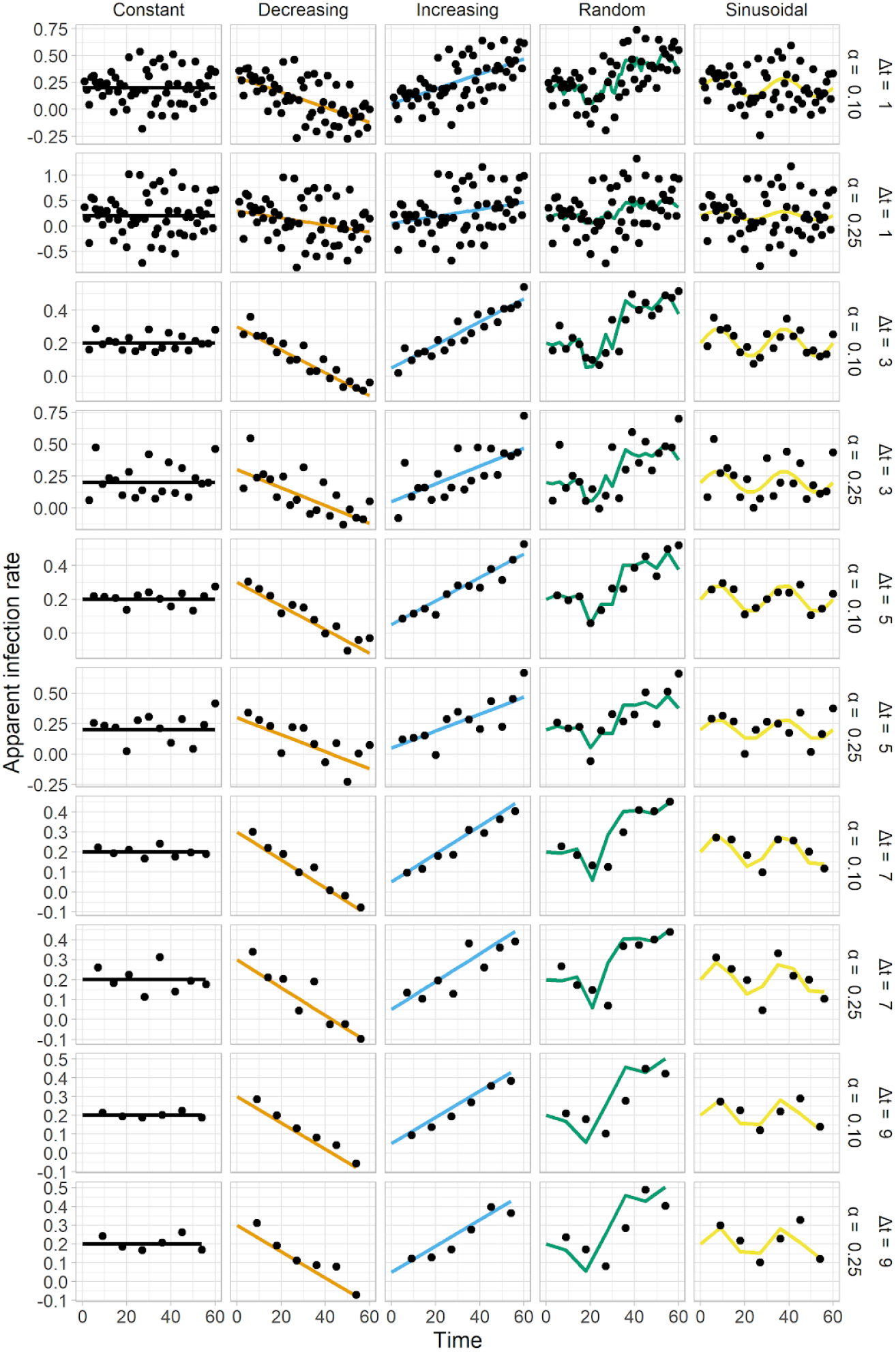
Calculation of the time-varying apparent infection rate, based on the rearrangement of the logistic equation, in synthetic disease progress curves simulated by a logistic model with five predefined patterns of time-varying apparent infection rate *r*(*t*) (constant, decreasing, increasing, random and sinusoidal), five intervals in time units (Δ*t* = 1, 3, 5, 7 or 9) and two levels of noise (α = 0.1 or 0.25) assigned to disease intensity data. Solid dots and triangles represent the calculated *r* for the respective level of noise and solid lines represents simulated values of *r*(*t*).

**Fig. S4.**
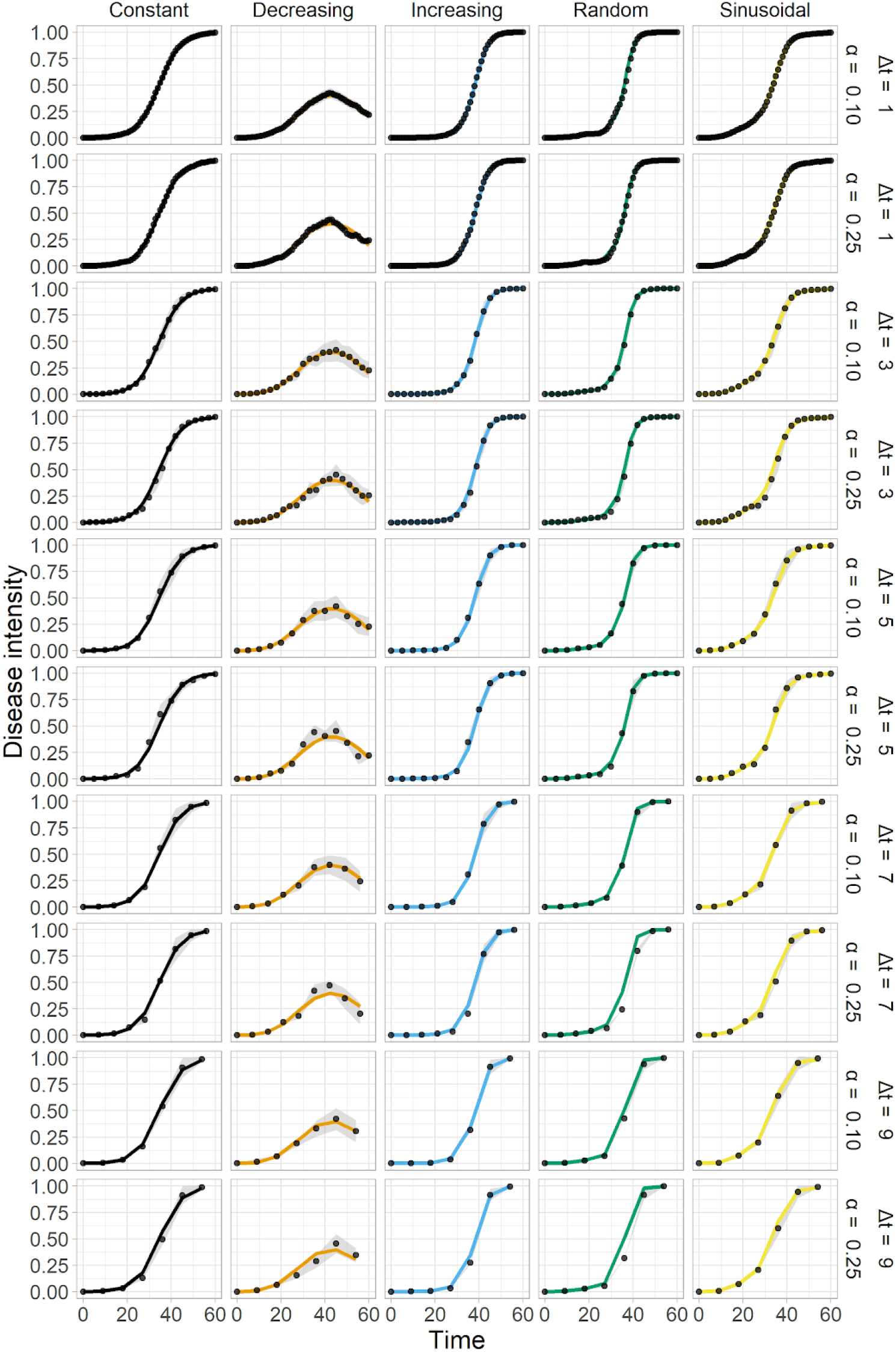
Disease progress curves of a disease intensity for a hypothetical polycyclic plant disease by a logistic model (solid line) and estimates of disease intensity by a particle filter approach for five patterns of time-varying apparent infection rate *r*(*t*) (constant, decreasing, increasing, random and sinusoidal), five intervals in time units (Δ*t* = 1, 3, 5, 7 or 9) and two levels of noise (α = 0.1 or 0.25) assigned to disease intensity data. Dots represent the state estimation by the SIR-PF and shaded area represent the the 99% confidence interval.

